# Scm^6^A: A fast and low-cost method for quantifying m^6^A modifications at the single-cell level

**DOI:** 10.1101/2023.12.14.571511

**Authors:** Yueqi Li, Jingyi Li, Wenxing Li, Shuaiyi Liang, Wudi Wei, Jiemei Chu, Jingzhen Lai, Yao Lin, Hubin Chen, Jinming Su, Xiaopeng Hu, Gang Wang, Jun Meng, Junjun Jiang, Li Ye, Sanqi An

**Author notes:** Corresponding author: (An S). Equal contribution (Yueqi Li, Jingyi Li, Wenxing Li, Shuaiyi Liang, Wudi Wei).

## Abstract

It is widely accepted that m^6^A exhibits significant intercellular specificity, which poses challenges for its detection using existing m^6^A quantitative methods. In this study, we introduce Scm^6^A, a machine learning-based approach for single-cell m^6^A quantification. Scm^6^A leverages input features derived from the expression levels of m^6^A *trans* regulators and *cis* sequence features, and found that Scm^6^A offers remarkable prediction efficiency and reliability. To further validate the robustness and precision of Scm^6^A, we applied a winscore-based m^6^A calculation method to conduct m^6^A-seq analysis on CD4^+^ and CD8^+^ T-cells isolated through magnetic-activated cell sorting (MACS). Subsequently, we employed Scm^6^A for analysis on the same samples. Notably, the m^6^A levels calculated by Scm^6^A exhibited a significant positive correlation with m^6^A quantified through m^6^A-seq in different cells isolated by MACS, providing compelling evidence for Scm^6^A’s reliability. We also used the scm^6^A-seq method to validate the reliability of our approach. Additionally, we performed single-cell level m^6^A analysis on lung cancer tissues as well as blood samples from COVID-19 patients, and demonstrated the landscape and regulatory mechanisms of m^6^A in different T-cell subtypes from these diseases. In summary, our work has yielded a novel, dependable, and accurate method for single-cell m^6^A detection. We are confident that Scm^6^A will have broad applications in the realm of m^6^A-related research.

## Introduction

As the most widespread epigenetic modification in mRNA, m^6^A plays pivotal roles in gene expression regulation and is intricately linked to physiological processes in various diseases[1–7]. Among its multifaceted regulatory functions, m^6^A governs T-cell differentiation and influences immune-related gene expression, garnering substantial attention[8]. To gain deeper insights into the role of m^6^A in biological progresses, it becomes imperative to discern transcriptome-wide m^6^A levels and sites within individual cells. For instance, the presence of a multitude of immune cell and T-cell subtypes[8–11] poses a formidable challenge, as current m^6^A detection methods designed for bulk cell populations fall short in characterizing m^6^A levels and sites at the single-cell level.

It is well-established that cell type-specific m^6^A levels and *de novo* m^6^A deposition are jointly regulated by *trans*-acting regulators and *cis*-regulatory elements[12]. In theory, leveraging information on these *trans*-acting regulators and *cis*-regulatory elements as input enables the prediction of m^6^A at the single-cell level using computational methods. Machine learning and other computational approaches have found extensive application in the analysis of diverse omics data, significantly advancing our understanding of biology [13–17]. In theory, machine learning holds the promise of predicting RNA methylation levels at the single-cell level. In our prior research, we developed a computational framework to systematically identify comprehensive *trans* regulators of m^6^A and performed experiments to verify the reliability of these *trans* regulators[18]. Additionally, a reliable regulatory network from *trans* regulators to m^6^A sites was constructed. Furthermore, we identified cell-specific m^6^A *cis*-regulatory motifs[18]. Machine learning, as a potent predictive tool, has been extensively employed in forecasting gene expression, DNA methylation, and alternative splicing, leveraging multiple biological features with impressive accuracy[19–22]. In fact, Xue et al. highlighted the challenges tied to the experimental detection of RNA m^6^A. To address this, they investigated the possibility of using computational methods to predict RNA methylation status based on gene expression data. Employing methods such as Support Vector Machine (SVM) and Random Forests (RF), Xue et al. determined that gene expression data can indeed act as a reliable predictor for m^6^A methylation status[23]. Their findings have convinced us of the viability of predicting single-cell m^6^A using machine learning methods grounded in gene expression level. Herein, we attempted to develop a single-cell level m^6^A calculation method through a machine learning method.

In this study, we leveraged comprehensive information on *trans*-regulatory elements of m^6^A and *cis*-elements, including motif and sequence data, to create a machine learning-based quantitative method for single-cell m^6^A analysis, which we named Scm^6^A (Single-cell m^6^A Analysis; available at https://github.com/Ansanqi/Scm6A). We applied multiple machine learning techniques to establish the association between *trans*-regulators and m^6^A, integrating *cis* sequence features and single-cell m^6^A levels. Subsequently, Scm^6^A was established with substantial predictive power to predict the level of m^6^A in individual cells. After that, we applied Scm^6^A to single-cell RNA-seq data from peripheral blood mononuclear cells (PBMCs) and calculated the m^6^A levels in CD4^+^ and CD8^+^ T-cell types. To validate the accuracy and reliability of Scm^6^A, we also performed m^6^A-seq on CD4^+^ and CD8^+^ T-cells, isolated via magnetic-activated cell sorting (MACS), from the same donor. Our findings underscored the precision and dependability of Scm^6^A in discerning single-cell m^6^A levels, in comparison to m^6^A-seq results derived from MACS-isolated cell populations. Subsequently, we extended our analysis to investigate single-cell m^6^A profiles in lung cancer scRNA-seq data using Scm^6^A and demonstrated that the m^6^A profiles are highly heterogeneous at the single-cell level in different subtypes of T-cells in lung cancer. We also applied our model to single-cell dataset of COVID-19[24], and demonstrated good performance in classifying T-cells and B-cells. Furthermore, we compared our Scm^6^A with the experimental method scm^6^A-seq developed by the Yang et al.[25], Scm^6^A not only performed well in mouse cells but also exhibited a significant correlation with experimental sequencing results.

## Results

### Random Forest Outperforms Other ML Models in Single-Cell m^6^A Calculation

Figure 1A illustrates the workflow of our study. Initially, we collected the gene expression data of 593 reliable m^6^A regulators and conserved sequence features associated with m^6^A, which we validated before[18]. To establish a precise single-cell m^6^A calculation model, we evaluated five machine learning regression models, including Random Forest (RF), K-NearestNeighbor (KNN), Support Vector Regression (SVR) with the poly kernel, Linear Regression (LR) and Linear Support Vector Regression (LinearSVR), all optimal parameters of the above models were obtained by grid search based on these *trans* and *cis* data.

**Figure 1.**
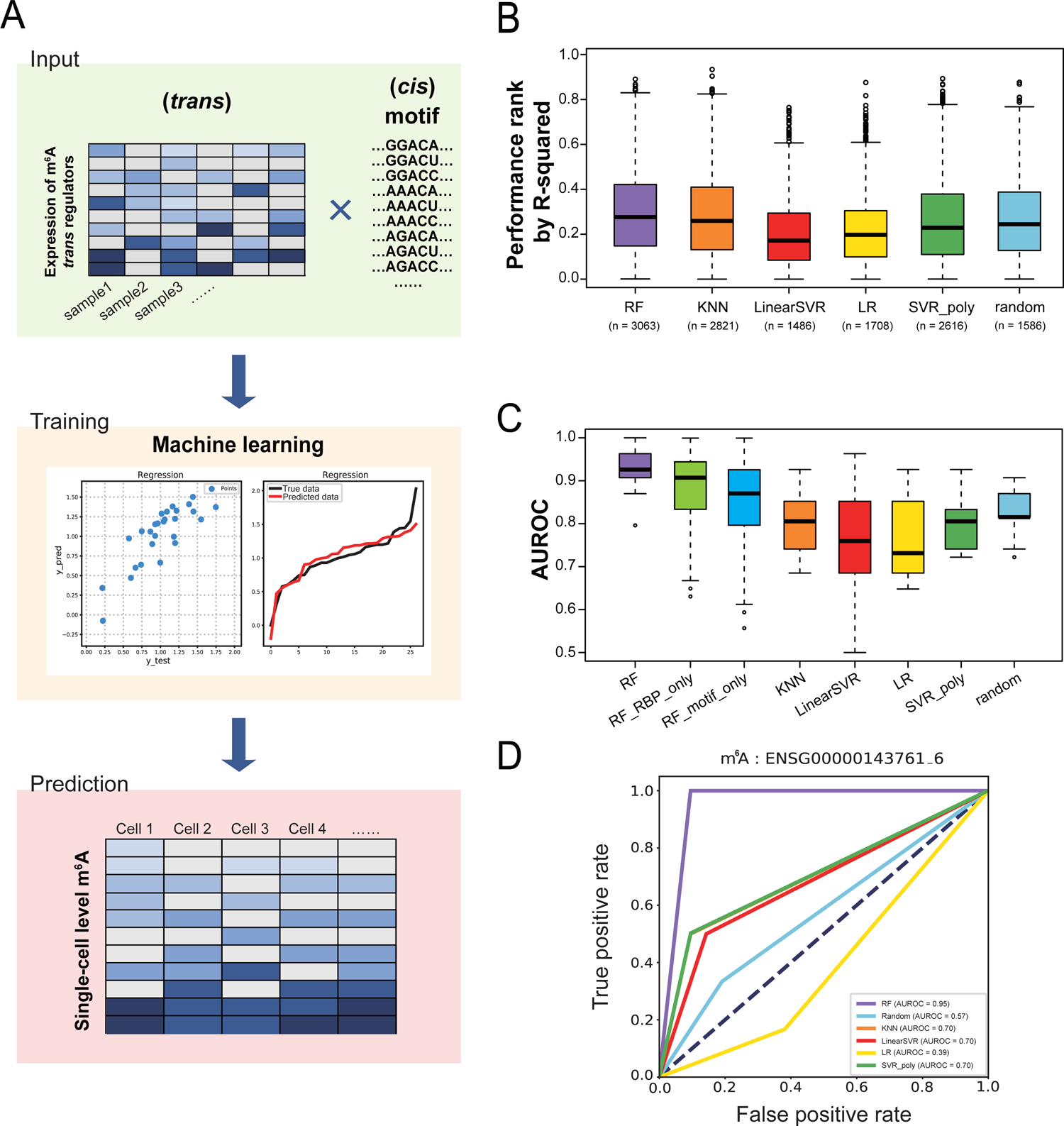
Single-cell m^6^A prediction models evaluation. A. Algorithmic framework for single-cell m^6^A calculation. B. Boxplot comparison rank of different models by R-squared in the test set. C. Area under the receiver operating characteristic (AUROC) for each method. D. ROC curves and AUC values for the different models in prediction of m^6^A sites in EMSG00000143761.

The coefficient of determination (R^2^), commonly employed to gauge the performance of regression-based machine learning models, was utilized as an evaluative parameter for assessing the proximity of data points to the fitted line. Notably, our analysis revealed that the RF model consistently outperformed the other machine learning models, displaying higher R^2^ values (Figure 1B), indicating that the RF model is the most suitable for single-cell m^6^A prediction. Additionally, the correlation analysis between the predicted m^6^A levels and true m^6^A levels demonstrated superior reliability of the RF model-based m^6^A calculation method compared to other models (**Figure S1A**). By defining the difference between the predicted value and the actual value as binary variables (See the Methods section for details), we performed receiver operating characteristic (ROC) analysis on models constructed using the five machine learning methods, based on their testing accuracy. Our findings revealed that the performance of Scm^6^A based on the RF model achieved a median balanced accuracy of 0.91 across multiple tests on all m^6^A sites (Figure 1C), which was substantially higher than that of other classical machine learning methods. Moreover, we conducted a comparative analysis between random forest models constructed solely using *trans*-acting regulators and those built solely using *cis*-regulatory elements. The median balanced accuracy of both models was found to be lower than that of the model created by integrating both types of effectors. The other four models achieved median balanced accuracies ranging from 0.8 to 0.9, suggesting the regulatory network we constructed is reliable. Moreover, an example is shown in Figure 1D. The ROC curves of the five models demonstrated distinct prediction performances, with the RF model exhibiting the highest prediction efficiency. To further validate our findings, we conducted a randomized relabeling of samples and performed ROC analysis using the methodology described above, the AUROC and R^2^ values of the RF model significantly exceeded those generated by random permutations (**Figure S1B, C**), suggesting a significant level of accuracy that cannot be explained by random chance. Overall, we identified the best fitting single-cell m^6^A calculation method and named it Scm^6^A.

### Accuracy and reliability of Scm^6^A were further validated by m^6^A-seq from magnetic-activated cell sorting in human PBMCs

We performed single-cell RNA-seq analysis of PBMCs from four healthy participants and extracted gene expression data for 593 reliable m^6^A regulators[18] as *trans*-acting input, and 42 m^6^A conserved sequence information[18, 26] as *cis*-acting input to for Scm^6^A. Using Scm^6^A, we calculated the single-cell level m^6^A profiles in CD4^+^ and CD8^+^ T-cells (Figure 2A). Simultaneously, we used a winscore-based m^6^A calculation method[18] to perform m^6^A-seq analysis of CD4^+^ and CD8^+^ T-cells isolated by MACS. To control technical biases in the regulatory network of *trans* m^6^A regulators to m^6^A sites in the m^6^A-seq libraries, including variations in sequencing lengths and RNA fragmentation lengths, we merged continuous Scm^6^A calculation peaks within the same gene, as described before[18]. Due to the different window sizes with two different calculation methods cannot be used for comparison of precise m^6^A locations, we obtained 49 precisely matched m^6^A windows to compare the correlations of m^6^A levels within the same gene region (Figure 2B), and these m^6^A sites with same localization on the transcriptome were the best choice for validating the accuracy and reliability of Scm^6^A. As we expected, there was a significant correlation between the m^6^A levels predicted by Scm^6^A and quantified by m^6^A-seq from MACS (Figure 2C). Moreover, there was no significant correlation between Scm^6^A and m^6^A-seq analysis results generated by random permutation of these m^6^A sites (Figure 2D), indicating a significant accuracy not be explained by random chances.

**Figure 2.**
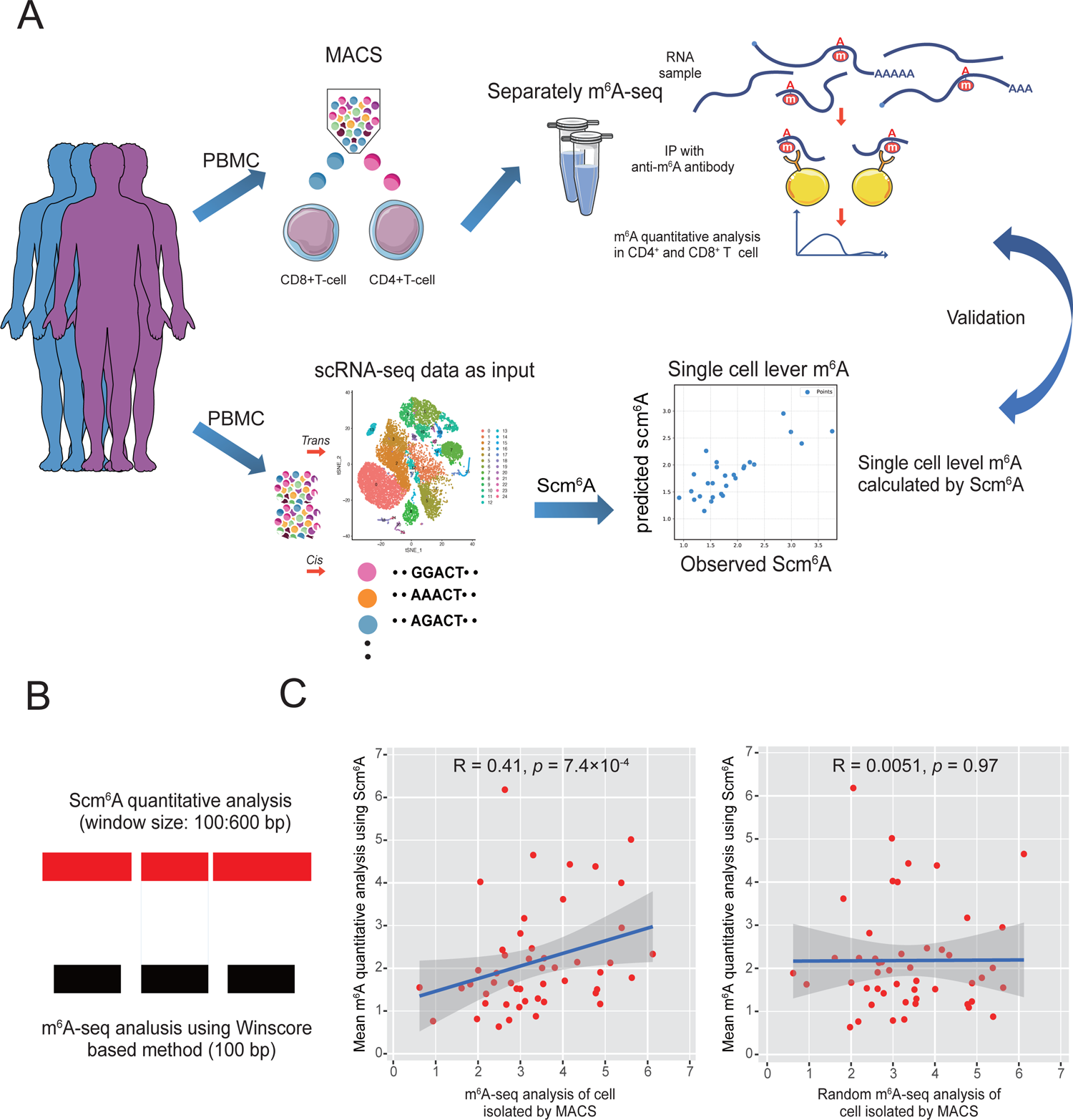
Verification of the reliability and accuracy of Scm^6^A. A. Schematic flow diagram showing the framework of validation of Scm^6^A. B. Schematic diagram of the distribution of m^6^A peak lengths calculated by Scm^6^A and the winscore-based method using m^6^A-seq. C. Correlation analysis between the Scm^6^A-calculated m^6^A levels and the winscore-based method-calculated m^6^A levels at the same m^6^A site. D. Correlation analysis between Scm^6^A calculated random m^6^A levels and winscore-based method calculated m^6^A levels at the same m^6^A site.

To further validate the accuracy and reliability of Scm^6^A, we conducted a comparative analysis between Scm^6^A and the single-cell m^6^A sequencing method recently published by Yang et al.[25], known as scm^6^A-seq. Given that scm^6^A-seq employs mouse cleavage-stage embryos cells as its experimental subject, we initially converted mouse gene IDs to their corresponding human gene IDs and subjected the gene expression data to standardized preprocessing. This step was carried out to enable an effective comparison with Scm^6^A. Despite variations in quantification methodologies, we harmonized the data through logarithmic transformations, ensuring that both Scm^6^A’s predictions and the m^6^A sequencing data provided in Yang et al.’s study could be juxtaposed for analysis.

Further correlation analysis results indicate a significant positive relationship (R=0.3, *p*=1×10^-74^) between the predictions generated by Scm^6^A and the experimentally measured m^6^A expression levels in scm^6^A-seq. This finding not only validates the efficacy of the Scm^6^A model in capturing underlying patterns within the data to a certain extent but also strongly reinforces the reliability of Scm^6^A’s predictive outcomes. We subsequently shuffled the order of the m^6^A sites and then performed a correlation analysis between the computational results of Scm^6^A and scm^6^A-seq. The correlation in the shuffled matrix was nearly absent (R=0, *p*=0.58) (**Figure S2A**). In summary, Scm^6^A proves to be a precise and dependable computational tool for single-cell m^6^A analysis. It’s cost-effectiveness, efficiency, and reliability make it a powerful tool with significant potential, offering researchers a rapid and effective means to explore the epigenetic features of cells.

### Identification of the potential role and landscape of m^6^A in CD4^+^ and CD8^+^ T-cells through m^6^A-seq

It is widely accepted that the differentiation of T-cell subtypes is associated with the expression of CD4 and CD8[27]. Transcriptional regulation plays a critical role in regulating the fate choice of CD4^+^/CD8^+^ T-cells[27]. Recently, some studies have reported that m^6^A has a broader impact on the dynamics of the RNA life cycle in T-cell differentiation by regulating crucial genes involved in T-cell differentiation[28].

Typically, a combination analysis of m^6^A-seq and RNA-seq is used to identify the potential role and mechanism of m^6^A-regulated genes in biological processes. To further validate the reliability of Scm^6^A, we performed m^6^A feature analysis using m^6^A-seq data from MACS and RNA-seq analysis to identify the potential differences in the role and landscape of m^6^A in CD4^+^ and CD8^+^ T-cells. As shown in Figure 3A, the m^6^A peaks of CD4^+^ T-cells tended to be enriched near stop codons, while the m^6^A peaks of CD8^+^ T-cells were enriched in coding regions and start codons, suggesting that the different T-cell types may have different m^6^A regulators controlling the m^6^A-mediated gene expression. We also checked the motif enrichment of CD4^+^ and CD8^+^ T-cells and found that m^6^A peaks in CD4^+^ T-cells more tended to be enriched in the GGACU motif. To be more specific, the *P*-value for motif enrichment analysis in CD4^+^ T-cells ranged from 1×10^-319^ to 1× 10^-407^, while the *P*-value in CD8^+^ T-cells ranged from 1×10^-207^ to 1×10^-282^(Figure 3B). Then, we performed different m^6^A analyses and differential expression analyses using m^6^A-seq and input data as RNA-seq data, as we reported before[18]. We found 2055 differentially expressed genes and 113 genes that contained different methylated m^6^A sites (278 m^6^A sites) (Figure 3C). The intersection of these two sets contains 62 genes (Figure 3D). We found that the gene expression levels of these genes were positively related to the m^6^A level of differentially methylated sites (Figure 3E**, F**). As expected, these potential m^6^A-regulated genes were enriched in pathways related to T-cell differentiation and cell differentiation, among others. (Figure 3G), suggesting m^6^A controls T-cell differentiation related gene expression.

**Figure 3.**
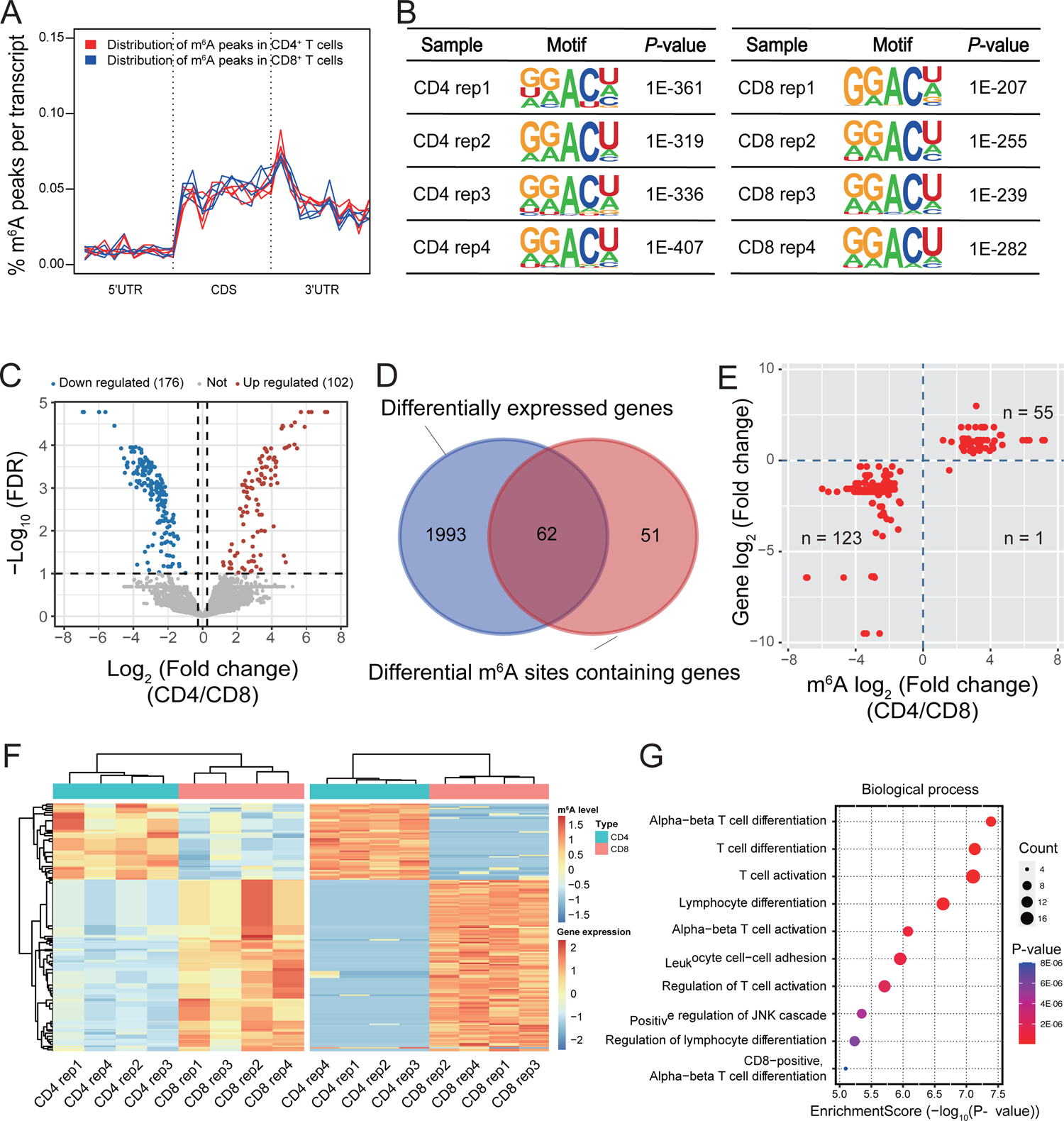
Bioinformatics analysis of m^6^A-sequencing results from MACS-isolated T-cells in human PBMC. **A**. The normalized distributions of m^6^A peaks across the 5′UTR, CDS and 3 ′UTR for CD4^+^ and CD8^+^ T-cells from four samples. **B.** Representative motifs for CD4^+^ and CD8^+^ T-cells in four samples. **C.** Volcano plots showing different m^6^A (FoldChange>1.2, FDR<0.1). Dots in red and blue indicate high and low expression of m^6^A and genes, respectively. **D.** Venn diagram of DEGs and genes where different m^6^A modifications are located. **E.** Scatter plot of log_2_FoldChange of different m^6^A and corresponding DEGs. **F.** Cluster gene expression heatmap of differentially expressed genes (Right) and m^6^A level heatmap of different m^6^A modifications (Left). **G.** Gene ontology enrichment analysis of the intersection of DEGs and different m^6^A.

### Identification of the potential role and landscape of m^6^A in CD4^+^ and CD8^+^ T-cells through Scm^6^A

Furthermore, we tried to investigate whether the combination analysis of Scm^6^A and single-cell RNA-seq performed as well as the combination analysis of m^6^A-seq and RNA-seq from MACS. As shown in Figure 4A, motif enrichment analysis revealed that m^6^A peaks calculated by Scm^6^A exhibited a tendency to be enriched in the GGACU motif in CD4^+^ T-cells, consistent with the enriched motif of m^6^A peaks identified through m^6^A-seq analysis (Figure 3B). To comprehensively analyze the m^6^A landscape at a single-cell resolution, we performed unsupervised clustering analysis of single-cell level m^6^A in CD4^+^ T-cells and CD8^+^ T-cells identified by scRNA-seq. We observed two clusters of single-cell m^6^A profiles (Figure 4B), which were clearly separated according to the cell types. Moreover, we predicted the m^6^A levels of CD4^+^ and CD8^+^ T-cells from a single sample at single-cell resolution and used cluster heatmaps to visualize the within-group similarity of the same cell type and the heterogeneity between groups of different cell types (Figure 4C), the genes containing these m^6^A modifications were enriched in T-cell differentiation and cell differentiation (Figure 4D). We also found that the gene expression levels of the m^6^A-deposited genes were positively related to the m^6^A of differentially methylated sites (Figure 4E), consistent with the results obtained from m^6^A-seq analysis using MACS-isolated cells (Figure 3F**, G**). These findings further underscore the reliability of Scm^6^A as a method for single-cell level m^6^A analysis.

**Figure 4.**
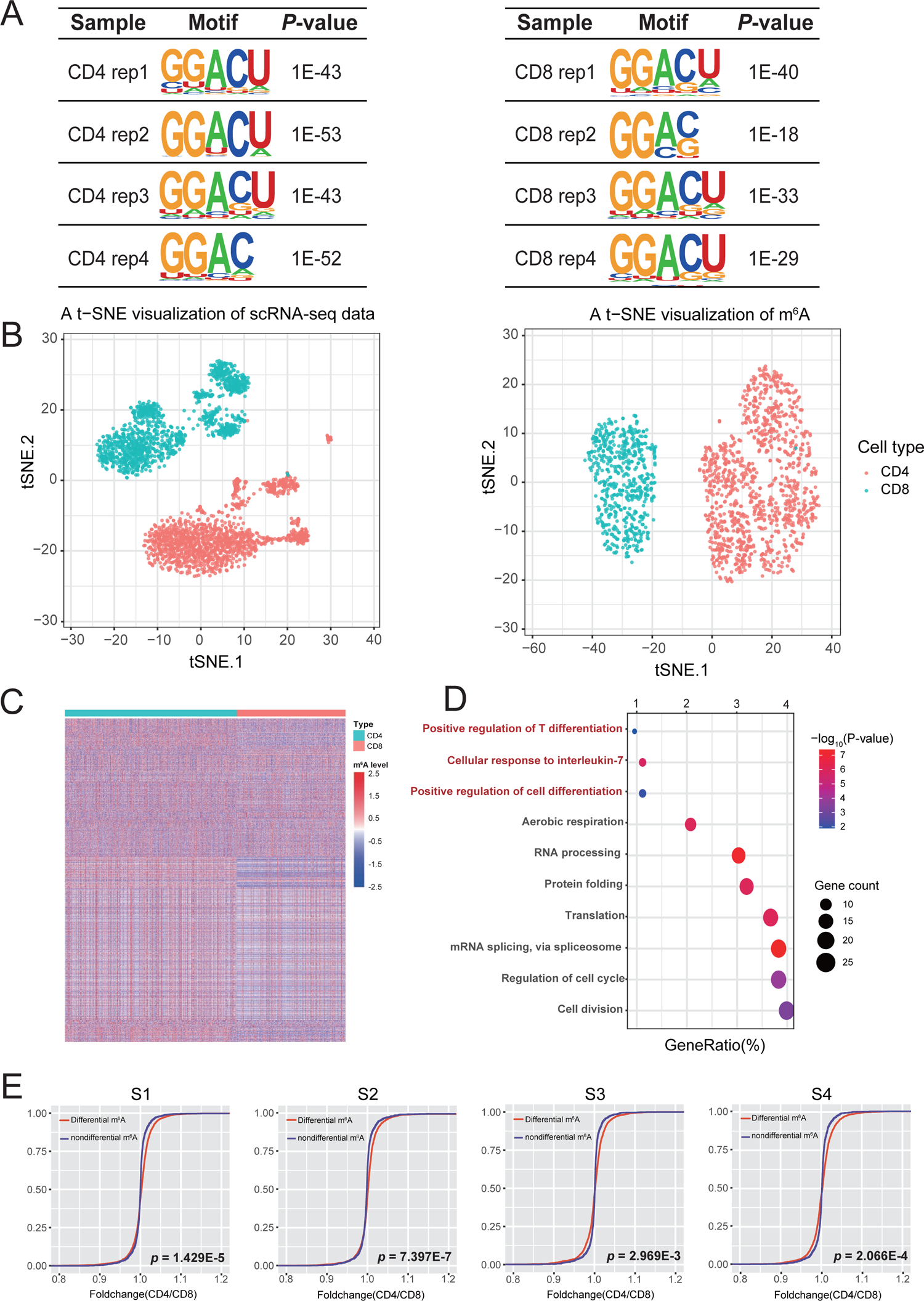
Bioinformatics analysis of Scm^6^A analysis results in human PBMC. **A**. Representative motifs for CD4^+^ and CD8^+^ T-cells in four samples. **B.** t-SNE maps of scRNA-seq data and m^6^A predicted by machine learning models. **C.** Cluster heatmap of differentially expressed m^6^A (FoldChange>1.2, FDR<0.1) predicted by machine learning models. **D.** Gene ontology enrichment analysis of the intersection of DEGs and Scm6A predicted different m^6^A located genes. **E.** Cumulative distribution function plots of different m^6^A and nondifferent m^6^A.

### Application of Scm^6^A in different lung cancer subtypes revealed the potential role and regulators of m^6^A in exhausted CD8^+^ T-cells

The differentiation of exhausted CD8^+^ T-cells leads to attenuated effector function of cytotoxic CD8^+^ T-cells, resulting in their inability to control tumor progression during the advanced stage [29]. Furthermore, it has been reported that m^6^A plays a crucial role in regulating T-cell homeostasis[8]. However, our current understanding of the different m^6^A profiles in distinct subtypes of T-cells and its role in exhausted CD8^+^ T-cells is limited[8].

Herein, we employed Scm^6^A to further explore the m^6^A landscape of exhausted CD8^+^ T-cells and other cell types across different lung cancer types, including lung squamous carcinoma (LUSC) (Figure 5A) and non-small cell lung cancer (NSCLC) (Figure 5B). Our findings revealed differences in the m^6^A profiles and molecular features of exhausted CD8^+^ T-cells compared to other T-cell subtypes in both LUSC and NSCLC (Figure 5A-C, **Figure S2B**). Interestingly, exhausted CD8^+^ T-cells (CD8_EM) -related m^6^A were enriched in IL-7 pathway, which is associated T-cell homeostasis (Figure 5D). By investigating the regulatory network we constructed (Figure 1A), it became evident that these exhausted CD8_EM-related m^6^A levels are associated with 19 m^6^A regulators, including METTL3, METTL14, and HMGB1 etc. (Figure 5E). Moreover, we observed a significant positive correlation between the expression levels of these 19 regulators and the m^6^A levels of CD8_EM-related m^6^A (Figure 5E). Therefore, we concluded that CD8_EM-related m^6^A regulators mediated m^6^A may regulate T-cell homeostasis through targeting IL-7. Consistent with this result, Hua-Bing et al. also found METTL3-mediated m^6^A controls T-cell homeostasis and differentiation by targeting IL-7, proving the reliability of our analysis results[8]. Notably, HMGB1, acting as a pivotal node in CD8_EM-related m^6^A regulation (Figure 5E), has previously been reported to influence the infiltration of CD8^+^ T-cells in NSCLC[30], further supporting the reliability of our conclusions.

**Figure 5.**
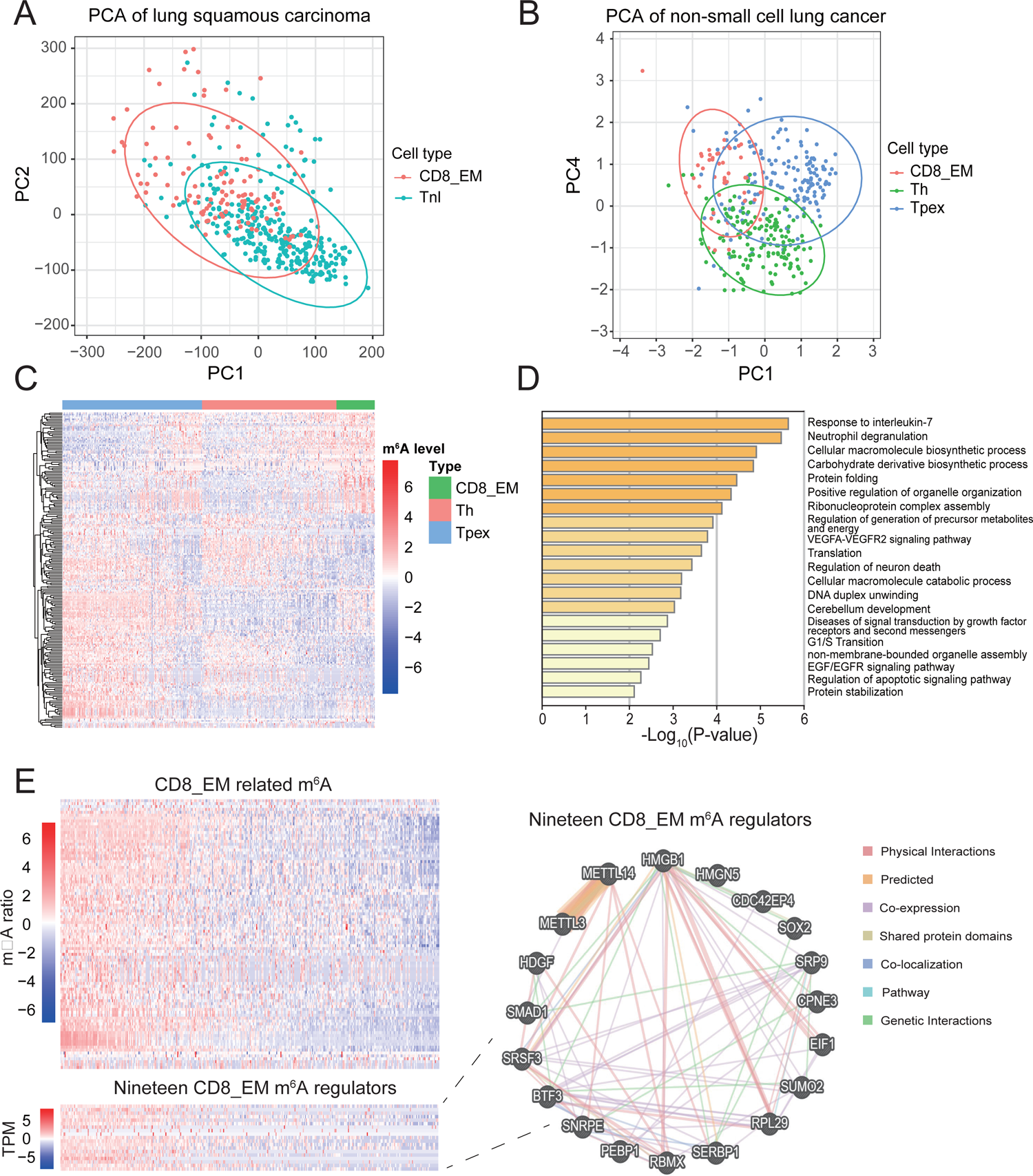
Using Scm^6^A to explore the single-cell level m^6^A from scRNA-seq data of lung cancer patients. **A.** and **B**. Principal component analysis (PCA) of single-cell m^6^A predicted by Scm^6^A. Each color represents a cell. The ellipses around the group mean represent the confidence regions. **C.** Heatmap of different single-cell m^6^A of NSCLC. **D.** Gene ontology enrichment analysis of the CD8_EM related m^6^A. **E.** Heatmaps representing the m^6^A ratios of the m^6^A peaks within CD8_EM related m^6^A (upper panel) and the gene expressions of the 19 CD8_EM related m^6^A regulators (lower panel) that significantly correlated with the m^6^A indexes of CD8_EM related m^6^A. These cells are sorted according to the m^6^A indexes of CD8_EM related m^6^A. Right panel shows Gene MANIA interaction network of 19 CD8_EM related m^6^A regulators with Physical Interactions, Predicted, co-expression, Shared protein domains, co-localization, Genetic Interactions and pathway. Dim grey nodes represent query regulators lists.

The dysregulation of immune responses in COVID-19 patients has emerged as a primary factor affecting symptoms and mortality rates[31, 32]. Consequently, the investigation of relevant immune cells has become a focal point in combatting this disease[33, 34]. In this context, we endeavored to apply Scm^6^A to a COVID-19 dataset from the study by Yuan et al[24]. From this dataset, we randomly selected a COVID-19 positive patient sample that had undergone FACS selection to isolate CD3^+^ T-cells and CD19^+^ B-cells from fresh PBMC. Using Seurat, we imported and standardized the single-cell data, then input the standardized data into Scm^6^A for prediction. The resulting UMAP plot of m^6^A predictions clearly delineated the classification of T-cells and B-cells (**Figure S3A**), highlighting a marked divergence in m^6^A modification landscapes between these cell types. Subsequent Gene Ontology (GO) functional enrichment analysis of the genes associated with differential m^6^A revealed significant enrichment in entries such as “SARS-CoV Infections” and “Viral Infection Pathways,” aligning with expectations and affirming the accuracy of Scm^6^A’s predictions (**Figure S3B**). These findings contribute to the dissection of the immune response in COVID-19 patients at a single-cell m^6^A resolution, enabling a deeper exploration of the pathogenic mechanisms at play.

These results provide a fresh perspective on the comprehensive profiles of m^6^A and corresponding regulators in exhausted CD8^+^ T-cells and other T-cells subtypes. Obviously, Scm^6^A has broaden applications in the identification of subtypes of different cell types, it may redefine some novel cell types and further reveal the potential role of m^6^A in cellular differentiation. This opens up exciting possibilities for future research in this field.

## Discussion

It has been reported that there is high heterogeneity in the abundance of m^6^A across individual cells[35]. However, a reliable and convenient method for detecting m^6^A at the single-cell level is currently lacking. In this study, we developed Scm^6^A, a single-cell m^6^A detection method based on the *trans* and *cis* information that we previously constructed. We validated the accuracy of the Scm^6^A method using m^6^A-seq data from MACS-isolated cell types. Subsequently, we applied Scm^6^A to single-cell RNA-seq data from lung cancer. Our analysis revealed that m^6^A is highly variable across different T-cells in lung cancer tissue. It also provides new ideas for the development of new single-cell calculation methods for other RNA methylation modifications, such as 5-methylcytosine (m^5^C), 7-methylguanosine (m^7^G), and N^1^-methyladenosine (m^1^A) methylation. Our attempts on the COVID-19 dataset have once again substantiated that Scm^6^A can effectively predict m^6^A expression levels and, in turn, distinguish between B-cells and T-cells. Moreover, we will further improve the application scope of Scm^6^A to analyze the single-cell level m^6^A in other species, including monkeys, and plant species, among others.

Recently, Matthew et al. developed a method named scDART-seq to identify transcriptome-wide YTH-binding m^6^A sites in single-cells by inducing APOBEC1-YTH expression[35]. However, this method can only be used to detect YTH-binding m^6^A sites, not all m^6^A sites. In addition, this method requires the expression of APOBEC1-YTH in targeted cells, which is not compatible with single-cell sequencing data. Moreover, scDART-seq can detect some false-positive m^6^A sites[35]. Due to the limitations of scDART-seq, there are no convenient and swift methods for identifying transcriptome-wide m^6^A sites and levels in individual cells with a low false discovery rate[36]. Comparing with scDART-seq, Scm^6^A offers a more accurate and convenient approach for quantifying m^6^A at the single-cell level. Therefore, Scm^6^A has more potential to be widely used in m^6^A-related research. Combining analysis of scRNA-seq with Scm^6^A also provides the pathway for researchers to investigate the detailed m^6^A regulation mechanisms at the single-cell level. Moreover, Scm^6^A displays high true positive rate with AUROC=0.91, suggesting that Scm^6^A can detect few false-positive m^6^A sites.

The role of m^6^A in T-cell differentiation and T-cell homeostasis has attracted much attention. As a key step of cancer immune evasion, there is a lack of research on the epigenetic mechanisms related to T-cell exhaustion[37], such as whether m^6^A is involved in regulating T-cell exhaustion. By using Scm^6^A, we found that the m^6^A profiles were distinct among these exhausted T-cells and other subtypes of T-cells, indicating that m^6^A plays an essential role in the progression of T-cell exhaustion. The impact of m^6^A on T-cell differentiation and activation deserves further discussion using our Scm^6^A method. Considering the discovery of pharmacological inhibition of METTL3[38], it is plausible that we may combat T-cell exhaustion by m^6^A-dependent gene regulation in the future. In fact, our method not only provides a single-cell m^6^A method for classifying T-cells but also extends to other cell type subpopulations, including B-cells, dendritic cells, and macrophages. We believe that Scm^6^A will be used to study the role of single-cell m^6^A in a variety of diseases and cell types.

Scm^6^A is based on a m^6^A-seq trained machine learning method and is antibody free. Even Scm^6^A is not a single-base resolution for every single transcript yet, we will incorporate new features of m^6^A identification using Nanopore Sequencing and miCLIP-m^6^A to Scm^6^A in the future[39–43]. We believed it will make Scm^6^A a more powerful tool for single-base resolution for every single transcript. Furthermore, we will try to collect more m^6^A sequencing data for training to improve the accuracy of Scm^6^A.

In summary, this study has provided a novel approach to the calculation of single-cell RNA methylation. Combined with other single-cell multiomics techniques, Scm^6^A will open up a new way for m^6^A research at the single-cell level which will be expected to significantly contribute to our understanding of the role of single-cell m^6^A in various biological processes.

## Methods

### Data preprocessing and machine learning algorithms

In accordance with our previous research, we collated and analyzed raw sequencing data from 104 m^6^A-seq libraries (IP and input) obtained from 25 unique cell lines, which were downloaded from the Sequence Read Archive (SRA, https://www.ncbi.nlm.nih.gov/sra). We identified comprehensive *trans* regulators in the m^6^A regulatory network and *cis* sequence features of the m^6^A site used these m^6^A-seq data [18]. Herein, we developed a single-cell level calculation method based on machine learning models using this network from the expression levels of the m^6^A regulators to the m^6^A level. As *cis*-acting regulatory sequence, position probability matrices of each m^6^A site were used as input to the model[44]. To solve the frequent NA values in m^6^A data, we first counted the percentage of missing values in each row and column and then deleted rows or columns with missing values greater than 10%. Following the data filtering process, the expression matrix consisting of 4162 m^6^A sites was obtained.

For the development of machine learning models, we established a one-to-one correspondence between the expression of m^6^A regulators and the m^6^A matrices to construct models. Meanwhile, considering the regulation of *cis*-acting elements to m^6^A, we included the RNA sequence of each m^6^A site converted into a matrix to the input information of the algorithm. In this study, a total of 5 machine learning algorithms, including RF, LR, KNN, LinearSVR, and SVR with poly kernel, were used to determine the most effective method. The scikit-learn toolkit version 1.0.2 was used to train these machine learning models. Out of the overall dataset, 70% was randomly allocated to serve as the training set for model development, while the remaining 30% was reserved for the test set to validate the model’s performance. The best parameters were selected through grid search and fivefold cross-validation.

### m^6^A-seq data processing

We established m^6^A-seq libraries (IP and input) of CD4^+^ T-cells and CD8^+^ T-cells isolated from the same individuals using MACS, and obtained four different sets of samples (Review Link: https://dataview.ncbi.nlm.nih.gov/object/PRJNA890754?reviewer=psjfcrhf1dk78idcnbo5a3msup). The reads generated by m^6^A-seq were mapped to the hg38 human reference genome using HISTA2 with default values for parameters (version 2.1.0)[45]. We calculated the input libraries’ TPM of Ensembl annotated genes using stringtie and then performed quantile normalization across all samples[46].

To accurately identify the m^6^A sites in CD4^+^ T-cells and CD8^+^ T-cells, we improved the winscore previously published by Dominissini et al.[36]. We determined the sliding window with a window fraction (enrichment fraction)>2 in the sample as the m^6^A peak. Since low expression windows may be accompanied by technical problems with unreliable winscore, we decided to adjust the windows with low RPKMs by adding 1 to the RPKM of each window in both IP and input libraries before winscore calculation. After identifying the m^6^A peaks across the samples, we merged consecutive m^6^A peaks within the same gene, and then divided the merged peaks with more than 5 consecutive sliding windows (300 bp) into multiple peaks, spanning no more than 5 sliding windows to eliminate the problem of possible false positives. After the above analyses, we finally obtained the m^6^A matrix of m^6^A levels in CD4^+^ T-cells and CD8^+^ T-cells.

### Calculation of single-cell RNA-seq

The “Seurat” package was used to analyze the single-cell sequencing data of CD4^+^ T-cells and CD8^+^ T-cells. Genes expressed in fewer than 3 cells and cells expressing fewer than 200 genes were excluded. Since cells with a high proportion of mitochondria-derived genes, a low number of detected genes, and a high proportion of unmapped or multi-mapped reads are often damaged or dying cells, which will affect the subsequent single-cell RNA-seq analysis, we performed quality control (QC) on the data to filter out the unqualified data. The parameters used were as follows: nFeature_RNA>200 & nFeature_RNA < 4000, nCount_RNA>200 & nCount_RNA < 20000, percent.mt < 25. To correct the variability of meaningful reads obtained by scRNA-seq across different cells, we normalized the expression using the “NormalizeData” function and calculated the top 1500 highly expressed variant genes by the “FindVariableFeatures” function.

To visualize and interpret the high-dimensional gene expression data, principal component analysis (PCA) and t-distributed stochastic neighbor embedding (tSNE) were used for visualizing scRNA-seq data of CD4^+^ T-cells and CD8^+^ T-cells. A total of 24 cell clusters were obtained after visualization. Using cellranger[47], we defined the cluster of cells with high CD4 expression as CD4^+^ T-cells and the cluster of cells with high CD8A and/or CD8B expression as CD8^+^ T-cells. After counting the marker gene expression levels of the four samples (S1-S4) and sorting them, we defined cluster 1 in the 24 clusters as CD8^+^ T-cells, and clusters 6, 9, 10, 11, 16, 18, 21 and 22 as CD4^+^ T-cells according to the marker gene expression level. To obtain differentially expressed genes, we used “DEseq2” to differentially analyze the genes of CD4^+^ and CD8^+^ T-cells obtained by scRNA-seq, and the parameters used were as follows: FDR<0.1 and FoldChange>1.2. After that, we used the “limma” package to differentially analyze the genes of m^6^A obtained by m^6^A-seq, and the parameters used were as follows: FDR<0.1. We downloaded lung cancer single-cell sequencing data from the GEO database as external data validation (GSE148071)[48], which contained 42 single-cell RNA sequencing data from tissues of stage III/IV NSCLC patients. We performed lung-cancer-related single-cell m^6^A analysis based on these scRNA-seq data. Through Scm^6^A, we calculated single-cell m^6^A levels using the expression levels of m^6^A regulators and position probability matrices of m^6^A sites as input from these scRNA-seq data.

### Human CD4^+^/CD8^+^ T-cell sorting

Human CD4^+^/CD8^+^ T-cells were obtained from PBMCs of adult donors in good health. The first step was to isolate PBMCs from whole blood by density gradient centrifugation (Ficoll 1.077 g/mL, Sigma-Aldrich, USA) and maintained in RPMI-1640 medium (Solarbio, China) supplemented with 15% FBS (Sigma-Aldrich, USA) and 1% penicillin/streptomycin (Solarbio, China). Second, magnetic activated cell sorting beads (MACS; Miltenyi Biotec) were used to isolate human CD4^+^/CD8^+^ T-cells from PBMCs. Briefly, PBMCs were bound to CD4 microbeads (20 µL of microbeads/10^7 cells) for 15 minutes at 4°C. After washing the cells with washing buffer, the cells were resuspended in 500 µL of washing buffer, passed through an LS column (Miltenyi) attached to a magnetic stand (Miltenyi) and washed three times. To elute targeted cells, the column was washed with buffer after being removed from the magnetic field. The approach was validated by MACS. The sorted targeted cells were used for single-cell PCR analysis and sequencing.

### GO analysis

To further investigate the mechanisms associated with the differences observed in different T-cells in healthy humans, we performed gene ontology (GO) functional enrichment analysis in DAVID (https://david.ncifcrf.gov/) by using the previously screened differentially expressed genes, and took the top 10 items of the biological process ranked in ascending order of FDR as the results[49].

### Motif and distribution of m^6^A peaks

By using the bed format file obtained earlier as the input file, Hypergeometric Optimization of Motif Enrichment (HOMER, http://homer.ucsd.edu/homer/) software was used for motif enrichment analysis. The distribution of m^6^A peaks was plotted on a mega gene with 10 bins in the 5’UTR, CDS, and 3’UTR regions, using the methods described in our previous paper[18].

### Statistical Analyses

Statistical analyses were performed using R version 4.0.2 (https://www.r-project.org/). Receiver operating characteristic curve (ROC) analysis and the area under the curve (AUC) were calculated using the pROC package to compare the efficacy of each model. We calculated ROC and AUC by randomly sampling the true and predicted values. Values were labeled as 1 if the difference between the predicted m^6^A value and the true value was no more than 0.5 and labeled as 0 if the difference fell outside the range. The evaluation indicators of the five-machine learning regressors did not conform to the normal distribution, so the quartile was used for statistics.

## Declarations

### Ethics approval and consent to participate

The studies involving human participants were reviewed and approved by Ethics and Human Subjects Committee of Guangxi Medical University (Ethical Review No.20210092). The patients/participants provided their written informed consent to participate in this study.

### Availability of data and materials

The raw data of the m^6^A-seq and single-cell RNA-seq data have been deposited in the Sequence Read Archive (SRA) database (https://www.ncbi.nlm.nih.gov/bioproject/PRJNA890754). The code associated with this research is available on GitHub at the following repository: https://github.com/Ansanqi/Scm6A

## Acknowledgments

This work was supported by the National Natural Science Foundation of China (82160389, 8210389), Guangxi Medical University Training Program for Distinguished Young Scholars to SQ.A., Guangxi Science and Technology Base and Talent Project (2022AC19006)

## CRediT authorship contribution statement

Yueqi Li: Formal analysis, Visualization, Writing – original draft, Writing – review & editing. Jingyi Li: Formal analysis, Writing – original draft. Wenxing Li: Formal analysis, Writing – original draft. Shuaiyi Liang: Formal analysis, Writing – original draft. Wudi Wei: Formal analysis, Writing – original draft. Jiemei Chu: Formal analysis. Jingzhen Lai: Formal analysis. Yao Lin: Visualization. Hubin Chen: Visualization. Jinming Su: Visualization. Xiaopeng Hu: Resources. Gang Wang: Resources. Jun Meng: Resources. Junjun Jiang: Resources, Conceptualization. Li Ye: Resources, Conceptualization. Sanqi An: Writing – original draft, Writing – review & editing, Conceptualization, Funding acquisition. All authors have read and approved the final manuscript.

---

**Figure S1.** A comparative analysis of m^6^A calculation methods based on the Random Forest model and other models A. Dots plot show the correlations between predicted m^6^A level and true m^6^A level. Boxplot show R-squared (B) and AUROC (C) of RF model use corresponding m^6^A regulators or random permutation to calculate single-cell m^6^A.

**Figure S2.** Application of Scm^6^A in datasets pertaining to mouse cleavage-stage embryos and lung cancer A. Scatterplots of correlation between Scm^6^A prediction results and the scm^6^A-seq sequencing outcomes (Left), and random correlation scatterplots of both (Right). B. Representative motifs for CD8_EM cells, Th and Tpex cells in NSCLC samples.

**Figure S3.** Application of Scm^6^A in COVID-19 data sets A. UMAP plot of m^6^A predicted by Scm^6^A. B. GO enrichment analysis of genes with differential m^6^A modifications.

## References

[1] Zhang Y, Chen W, Zheng X, Guo Y, Cao J, Zhang Y, et al. Regulatory role and mechanism of m(6)A RNA modification in human metabolic diseases. Mol Ther Oncolytics 2021; (22):52–63.

[2] Wang X, Lu X, Wang P, Chen Q, Xiong L, Tang M, et al. SRSF9 promotes colorectal cancer progression via stabilizing DSN1 mRNA in an m6A-related manner. Journal of Translational Medicine 2022; (20).

[3] Niu X, Yang Y, Ren Y, Zhou S, Mao Q, Wang Y. Crosstalk between m(6)A regulators and mRNA during cancer progression. Oncogene 2022; (41):4407–19.

[4] Meyer KD. HOW M(6)A MAKES ITS MARK. Nature Reviews Molecular Cell Biology 2022; (23):519-.

[5] Lee Y, Choe J, Park OH, Kim YK. Molecular Mechanisms Driving mRNA Degradation by m(6)A Modification. Trends in Genetics 2020; (36):177–88.

[6] Khan RIN, Malla WA. m(6)A modification of RNA and its role in cancer, with a special focus on lung cancer. Genomics 2021; (113):2860–9.

[7] An S, Xie Z, Liao Y, Jiang J, Dong W, Yin F, et al. Systematic analysis of clinical relevance and molecular characterization of m6A in COVID-19 patients. Genes & Diseases 2022; (9):1170–3.

[8] Li H-B, Tong J, Zhu S, Batista PJ, Duffy EE, Zhao J, et al. m(6)A mRNA methylation controls T cell homeostasis by targeting the IL-7/STAT5/SOCS pathways. Nature 2017; (548):338–42.

[9] Juarez I, Su S, Herbert Z, Teijaro J, Moulton VJFii. Splicing factor SRSF1 is essential for CD8 T cell function and host antigen-specific viral immunity. 2022; (13).

[10] Baharom F, Ramirez-Valdez RA, Khalilnezhad A, Khalilnezhad S, Dillon M, Hermans D, et al. Systemic vaccination induces CD8+ T cells and remodels the tumor microenvironment. 2022.

[11] Al Moussawy M, Abdelsamed HAJFiI. Non-cytotoxic functions of CD8 T cells:“repentance of a serial killer”. 2022; (13).

[12] Huang H, Weng H, Chen J. The Biogenesis and Precise Control of RNA m(6)A Methylation. Trends in Genetics 2020; (36):44–52.

[13] Yang Z, Yi W, Tao J, Liu X, Zhang MQ, Chen G, et al. HPVMD-C: a disease-based mutation database of human papillomavirus in China. Database (Oxford) 2022; (2022).

[14] Yang S, Wang Y, Chen Y, Dai Q. MASQC: Next Generation Sequencing Assists Third Generation Sequencing for Quality Control in N6-Methyladenine DNA Identification. Front Genet 2020; (11):269.

[15] Wang Y, Xu Y, Yang Z, Liu X, Dai Q. Using Recursive Feature Selection with Random Forest to Improve Protein Structural Class Prediction for Low-Similarity Sequences. Comput Math Methods Med 2021; (2021):5529389.

[16] Kong R, Xu X, Liu X, He P, Zhang MQ, Dai Q. 2SigFinder: the combined use of small-scale and large-scale statistical testing for genomic island detection from a single genome. Bmc Bioinformatics 2020; (21):159.

[17] Dai Q, Bao C, Hai Y, Ma S, Zhou T, Wang C, et al. MTGIpick allows robust identification of genomic islands from a single genome. Brief Bioinform 2018; (19):361–73.

[18] An S, Huang W, Huang X, Cun Y, Cheng W, Sun X, et al. Integrative network analysis identifies cell-specific trans regulators of m(6)A. Nucleic Acids Research 2020; (48):1715–29.

[19] Li J, Guo G. Deciphering single-cell transcriptional programs across species. Nature Genetics 2022.

[20] Levy JJ, Titus AJ, Petersen CL, Chen Y, Salas LA, Christensen BC. MethylNet: an automated and modular deep learning approach for DNA methylation analysis. Bmc Bioinformatics 2020; (21).

[21] He J, Sun M-a, Wang Z, Wang Q, Li Q, Xie H. Characterization and machine learning prediction of allele-specific DNA methylation. Genomics 2015; (106):331–9.

[22] Cai Q, He B, Zhang P, Zhao Z, Peng X, Zhang Y, et al. Exploration of predictive and prognostic alternative splicing signatures in lung adenocarcinoma using machine learning methods. Journal of Translational Medicine 2020; (18).

[23] Xue H, Wei Z, Chen K, Tang Y, Wu X, Su J, et al. Prediction of RNA Methylation Status From Gene Expression Data Using Classification and Regression Methods. Evol Bioinform Online 2020; (16):1176934320915707.

[24] Ren X, Wen W, Fan X, Hou W, Su B, Cai P, et al. COVID-19 immune features revealed by a large-scale single-cell transcriptome atlas. Cell 2021; (184):1895–913.e19.

[25] Yao H, Gao CC, Zhang D, Xu J, Song G, Fan X, et al. scm(6)A-seq reveals single-cell landscapes of the dynamic m(6)A during oocyte maturation and early embryonic development. Nat Commun 2023; (14):315.

[26] Linder B, Grozhik AV, Olarerin-George AO, Meydan C, Mason CE, Jaffrey SRJNm. Single-nucleotide-resolution mapping of m6A and m6Am throughout the transcriptome. 2015; (12):767–72.

[27] Ellmeier W, Haust L, Tschismarov R. Transcriptional control of CD4 and CD8 coreceptor expression during T cell development. Cellular and Molecular Life Sciences 2013; (70):4537–53.

[28] Furlan M, Galeota E, de Pretis S, Caselle M, Pelizzola M. m6A-Dependent RNA Dynamics in T Cell Differentiation. Genes 2019; (10).

[29] Giles JR, Ngiow SF, Manne S, Baxter AE, Khan O, Wang P, et al. Shared and distinct biological circuits in effector, memory and exhausted CD8(+) T cells revealed by temporal single-cell transcriptomics and epigenetics. Nat Immunol 2022.

[30] Gao Q, Wang S, Chen X, Cheng S, Zhang Z, Li F, et al. Cancer-cell-secreted CXCL11 promoted CD8(+) T cells infiltration through docetaxel-induced-release of HMGB1 in NSCLC. J Immunother Cancer 2019; (7):42.

[31] Lin Y, Li Y, Chen H, Meng J, Li J, Chu J, et al. Weighted gene co-expression network analysis revealed T cell differentiation associated with the age-related phenotypes in COVID-19 patients. BMC Med Genomics 2023; (16):59.

[32] Chen Y, Wang G, Li J, Xia L, Zhu L, Li W, et al. CASA: a comprehensive database resource for the COVID-19 Alternative Splicing Atlas. J Transl Med 2022; (20):473.

[33] An S, Xie Z, Liao Y, Jiang J, Dong W, Yin F, et al. Systematic analysis of clinical relevance and molecular characterization of m(6)A in COVID-19 patients. Genes Dis 2022; (9):1170–3.

[34] An S, Li Y, Lin Y, Chu J, Su J, Chen Q, et al. Genome-Wide Profiling Reveals Alternative Polyadenylation of Innate Immune-Related mRNA in Patients With COVID-19. Front Immunol 2021; (12):756288.

[35] Tegowski M, Flamand MN, Meyer KD. scDART-seq reveals distinct m(6)A signatures and mRNA methylation heterogeneity in single cells. Molecular cell 2022; (82):868-+.

[36] Dominissini D, Moshitch-Moshkovitz S, Salmon-Divon M, Amariglio N, Rechavi G. Transcriptome-wide mapping of N-6-methyladenosine by m(6)A-seq based on immunocapturing and massively parallel sequencing. Nature Protocols 2013; (8):176–89.

[37] Dolina JS, Van Braeckel-Budimir N, Thomas GD, Salek-Ardakani S. CD8(+) T Cell Exhaustion in Cancer. Front Immunol 2021; (12):715234.

[38] Yankova E, Blackaby W, Albertella M, Rak J, De Braekeleer E, Tsagkogeorga G, et al. Small-molecule inhibition of METTL3 as a strategy against myeloid leukaemia. Nature 2021; (593):597–601.

[39] Li JH, Liu S, Zhou H, Qu LH, Yang JH. starBase v2.0: decoding miRNA-ceRNA, miRNA-ncRNA and protein-RNA interaction networks from large-scale CLIP-Seq data. Nucleic Acids Res 2014; (42):D92–7.

[40] Shah A, Qian Y, Weyn-Vanhentenryck SM, Zhang C. CLIP Tool Kit (CTK): a flexible and robust pipeline to analyze CLIP sequencing data. Bioinformatics 2017; (33):566–7.

[41] Liu H, Begik O, Novoa EM. EpiNano: Detection of m(6)A RNA Modifications Using Oxford Nanopore Direct RNA Sequencing. Methods Mol Biol 2021; (2298):31–52.

[42] Parker MT, Knop K, Sherwood AV, Schurch NJ, Mackinnon K, Gould PD, et al. Nanopore direct RNA sequencing maps the complexity of Arabidopsis mRNA processing and m(6)A modification. Elife 2020; (9).

[43] Zhang Y, Huang D, Wei Z, Chen K. Primary sequence-assisted prediction of m(6)A RNA methylation sites from Oxford nanopore direct RNA sequencing data. Methods 2022; (203):62–9.

[44] Gao Z, Liu L, Ruan JH. Logo2PWM: a tool to convert sequence logo to position weight matrix. BMC Genomics 2017; (18).

[45] Kim D, Langmead B, Salzberg SL. HISAT: a fast spliced aligner with low memory requirements. Nature Methods 2015; (12):357–U121.

[46] Pertea M, Pertea GM, Antonescu CM, Chang T-C, Mendell JT, Salzberg SL. StringTie enables improved reconstruction of a transcriptome from RNA-seq reads. Nature Biotechnology 2015; (33):290-+. cellranger:https://support.10xgenomics.com/single-cell-dna/software/overview/welcome.

[47] Wu F, Fan J, He Y, Xiong A, Yu J, Li Y, et al. Single-cell profiling of tumor heterogeneity and the microenvironment in advanced non-small cell lung cancer. Nat Commun 2021; (12):2540.

[48] Huang DW, Sherman BT, Lempicki RA. Bioinformatics enrichment tools: paths toward the comprehensive functional analysis of large gene lists. Nucleic Acids Research 2009; (37):1–13.

