## Supplementary figures and images for "Scm^6^A: A fast and low-cost method for quantifying m^6^A modifications at the single-cell level"

### Supplementary Figure 1

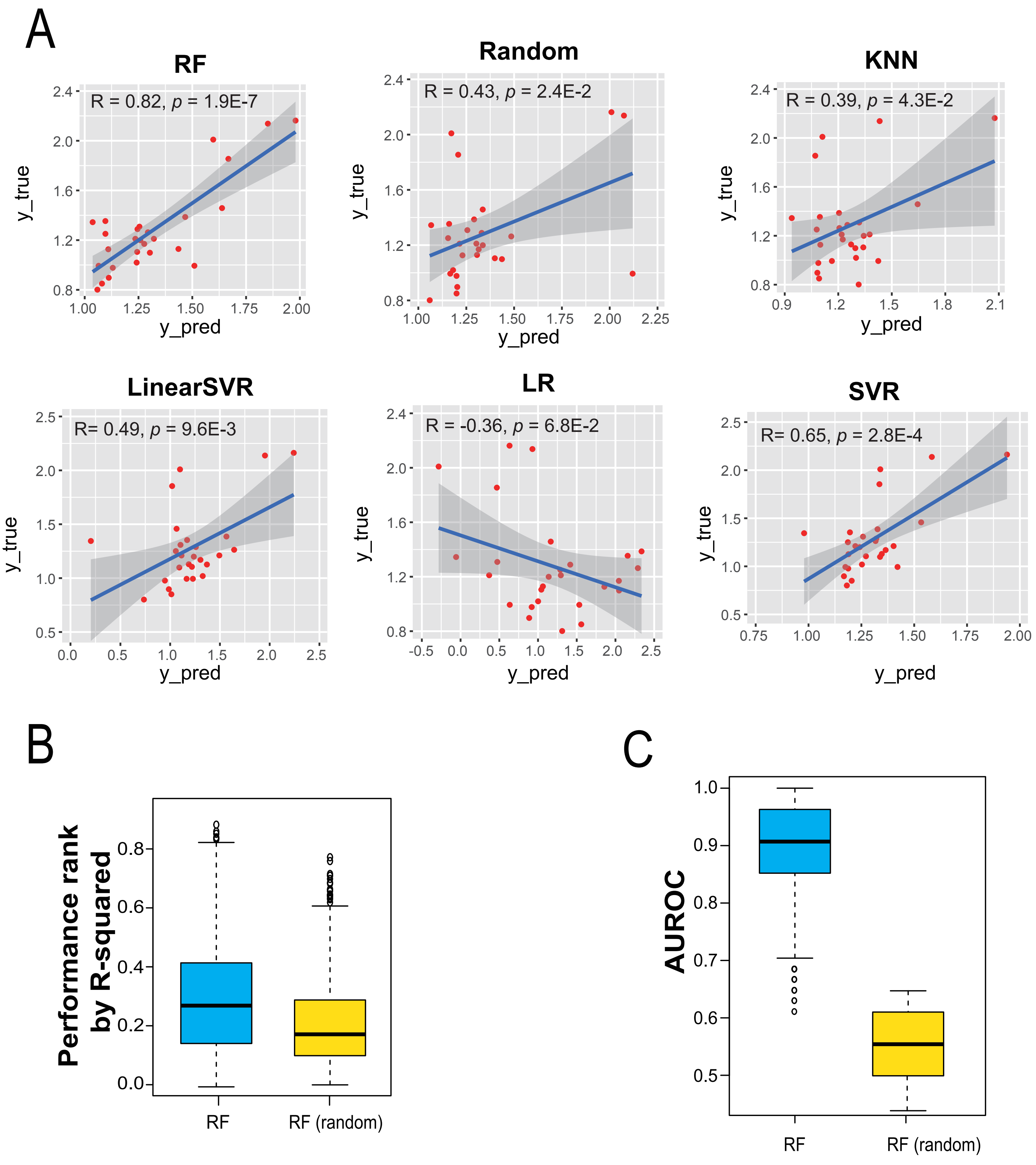

### Supplementary Figure 2

A

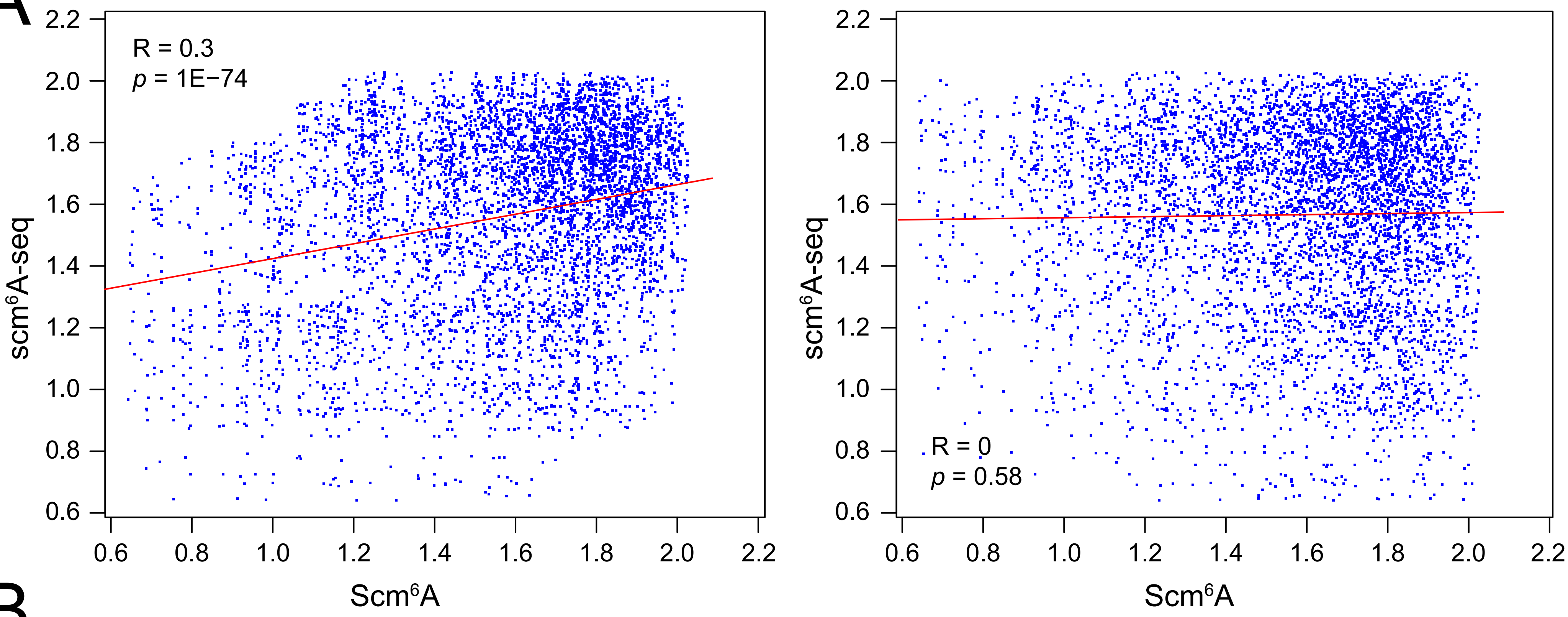


B

| **Cluster** | **Motif** | ***P*-value** |
|---|---|---|
| CD8_EM | 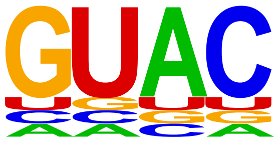 | 1E-3 |
| Th | 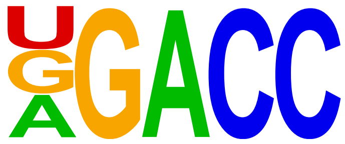 | 1E-7 |
| Tpex | 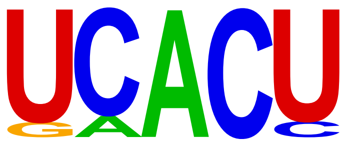 | 1E-10 |

### Supplementary Figure 3

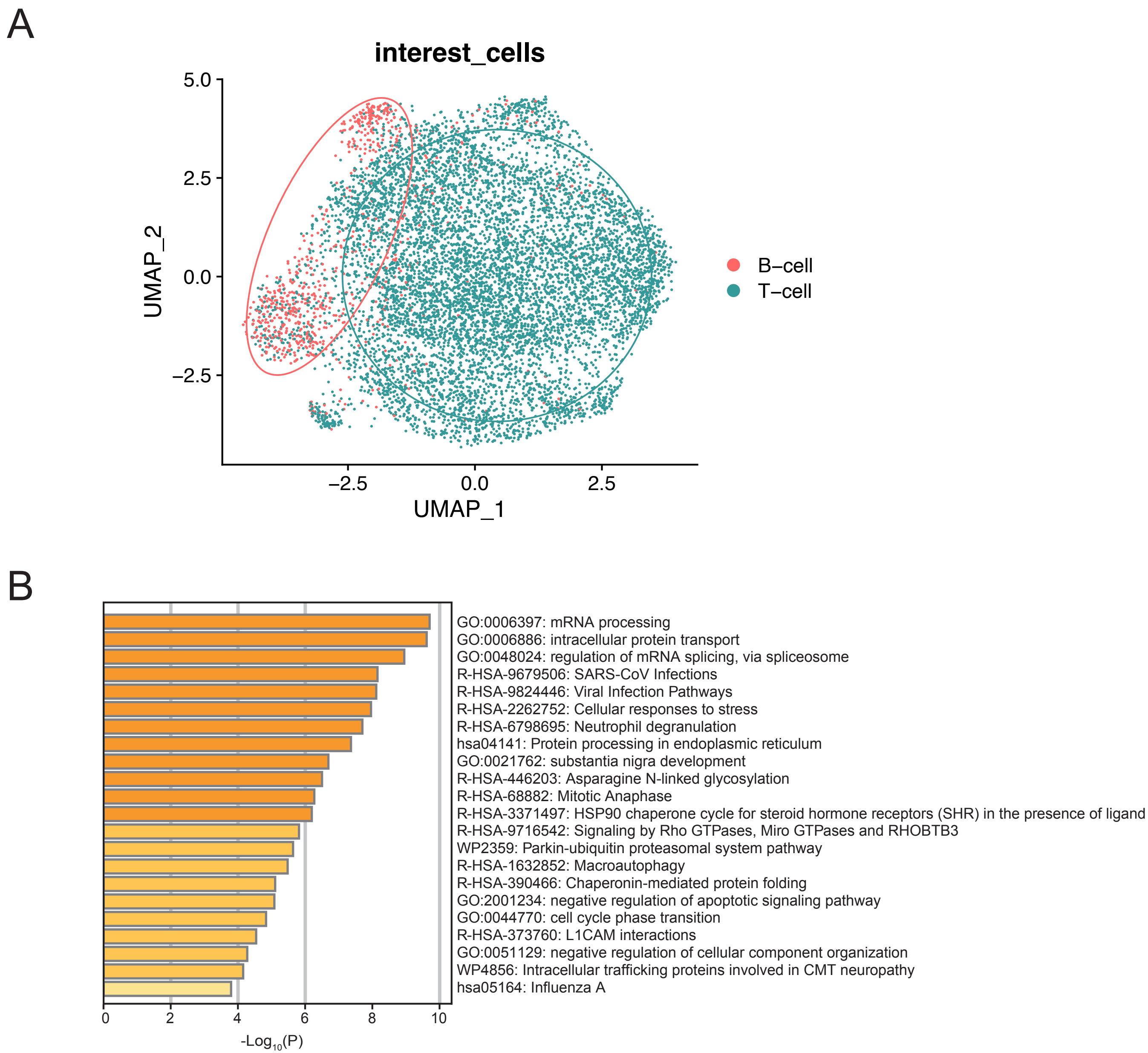
